# Metabolic engineering of narrow-leafed lupin for the production of enantiomerically pure (‒)-sparteine

**DOI:** 10.1101/2024.04.26.591273

**Authors:** Davide Mancinotti, Ting Yang, Fernando Geu-Flores

**Affiliations:** Section for Plant Biochemistry and Copenhagen Plant Science Centre, Department of Plant and Environmental Sciences, University of Copenhagen, Frederiksberg, Denmark

## Abstract

Sparteine is a plant-derived alkaloid widely known for its utility as chiral ligand in asymmetric synthesis^1-3^. However, its variable market price and availability have failed to meet the demand for a cheap and reliable product^4-6^. Sparteine is naturally synthesized by a sub-group of legume plants, which typically accumulate complex mixtures of closely related alkaloids. Here, we identified two enzymes from narrow-leafed lupin (NLL, *L. angustifolius*) that can sequentially oxidize (−)-sparteine to (+)-lupanine. The first enzyme is a cytochrome P450 monooxygenase belonging to family 71 (CYP71D189) and the second one is a short-chain dehydrogenase/reductase (SDR1). We also screened a non-GMO NLL mutant library and isolated a knockout in CYP71D189. The knockout displayed an altered metabolic profile where (−)-sparteine accounted for 96% of the alkaloid content in the seeds (GC-MS basis). The (−)-sparteine isolated from the mutant seeds was enantiomerically pure (99% *ee*). Apart from the altered alkaloid profile, the mutant did not have any noticeable phenotype. Our work demonstrates that (−)-sparteine is the precursor of most QAs in NLL and provides a convenient source of this valuable compound for academia and industry.

## Main

Asymmetric synthesis is the synthesis of chiral molecules using methods that favor the formation of a specific enantiomer/diastereomer. Asymmetric synthesis is particularly relevant for the pharmaceutical industry, as different enantiomers/diastereomers typically have different biological activities. One of the most versatile auxiliary molecules used in asymmetric synthesis is the chiral diamine sparteine. Specifically, complexes between sparteine and lithium have proven unrivaled for asymmetric deprotonations, substitutions, carbometalations, and directed *orto-*metalations^7 and references therein^. As chemists continue to find new uses for sparteine-lithium complexes, their versatility continues to grow, as exemplified by the development of assembly-line chiral synthesis^8, 9^. Originally, only (−)-sparteine was commercially available, enabling the asymmetric synthesis of just one of two possible products in each case. This led to the development of a synthetically accessible molecule with the expected functionality of (+)-sparteine, which became known as the (+)-sparteine surrogate^7^. Commercial (−)-sparteine remained cheap throughout the 2000s^10^; however, for reasons that remain unclear, it later became expensive and, at times, fully unavailable^4^. The fluctuations in price and availability motivated the development of a (−)-sparteine surrogate^11^ as well as an improved synthetic route for (−)-sparteine itself^12^. However, the multistep nature of these protocols (8 and 10 steps, respectively)^12^ has prevented their adoption by the wider community.

Sparteine belongs to the family of quinolizidine alkaloids (QAs), which are produced by leguminous plants of the larger genistoid clade^13^. Bicyclic, tricyclic, and tetracyclic QAs exist, with sparteine being the simplest tetracyclic one (see structure in Fig. 1). By and large, the most studied QA-producing plants are the lupins (*Lupinus* spp.), which typically produce complex mixtures of QAs sometimes including sparteine^14^. The enantiomeric purity of lupin-derived QAs is species-dependent. For example, the tetracyclic QA lupanine occurs in white lupin (*L. albus*) as a near-racemic mixture^15^, while the same QA in narrow-leafed lupin (NLL, *L. angustifolius*) seems to occur exclusively as the dextrorotatory form^16^ (Fig. 1a). Since the seeds of white lupin are consumed as snacks in Southern Europe upon alkaloid removal, chemists have developed methods for the chiral resolution of lupanine and its subsequent reduction to sparteine^17-19^. Interestingly, several wild North American lupins predominantly accumulate (−)-sparteine, including *L. babiger*^20^ and *L. montanus*^21^. Outside the *Lupinus* genus, few species are known to display similar characteristics, in particular, scotch broom (*Cytisus scoparius*, also known as *Sarothamnus scoparius*)^22, 23^, which appears to have been the source of commercial (−)-sparteine throughout the 1990s and 2000s^4^. The extent to which any of these (−)-sparteine-accumulating plants can become a reliable commercial source of the alkaloid remains unclear, as these are all wild species not optimized for production and likely prone to large seasonal variation.

**Figure 1.**
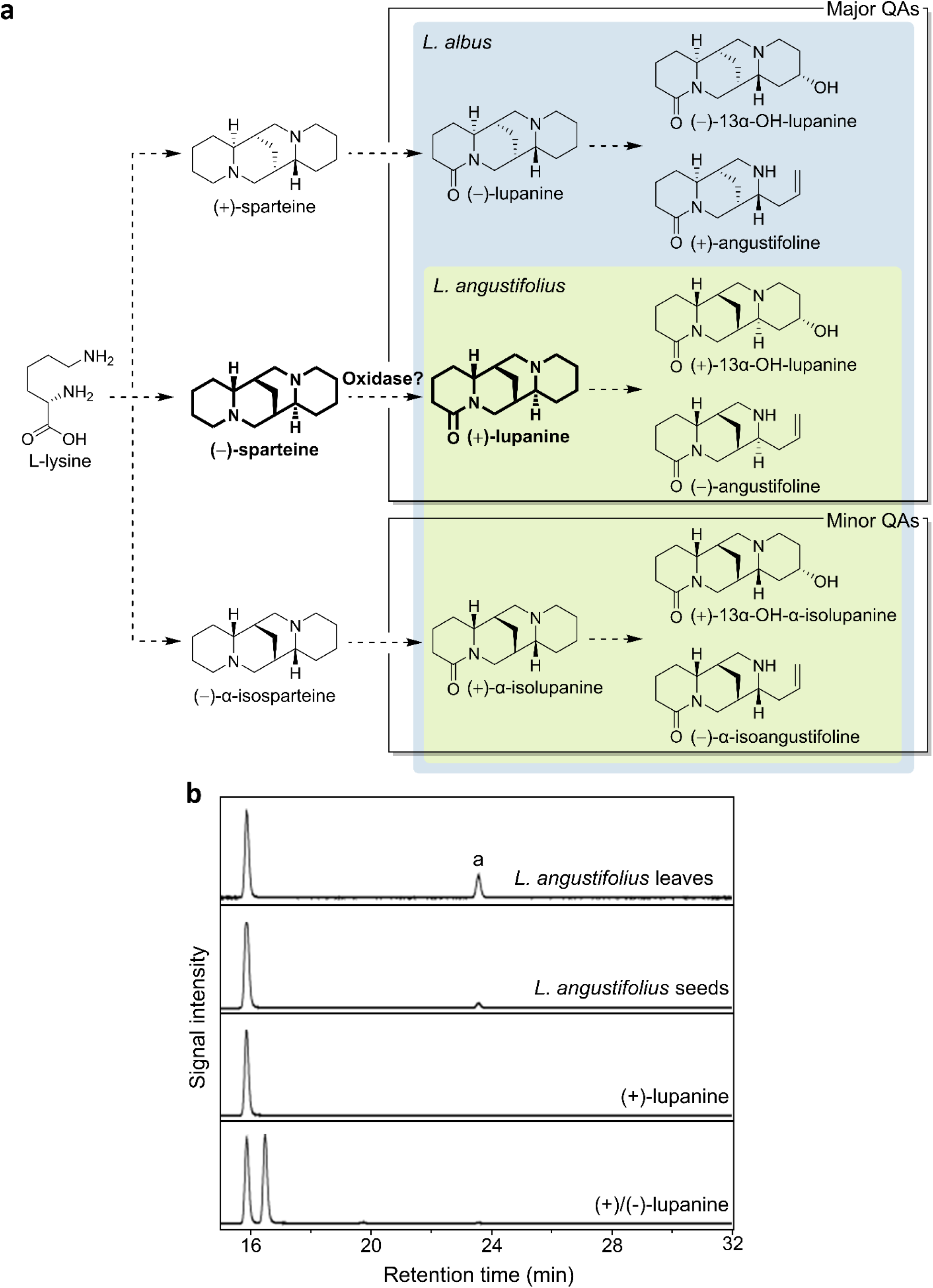
QA occurrence and biosynthesis in two common lupin crops, white lupin (*L. albus*) and narrow-leafed lupin (NLL, *L. angustifolius*). **a**. Overall biosynthetic proposal based on the backbone structure of the QAs found in each species. The proposed two-step enzymatic oxidation of (−)-sparteine to (+)-lupanine (in bold) is the subject of this study. In addition to the QAs shown above, white lupin also accumulates multiflorine and derivatives^34^. **b**. Chiral LC-MS analysis of NLL seed and leaf extracts, showing the enantiopurity of (+)-lupanine as well as small amounts of α-isolupanine (peak a). Traces are extracted ion chromatograms at *m/z* 249.20 ± 0.01 ([M+H]^+^ of lupanine).

Our group is currently studying the biosynthesis and transport of QAs in NLL, a cultivated lupin species grown in Europe and Australia for its protein-rich seeds. NLL does not normally accumulate (−)-sparteine but accumulates at least eight related QAs, three of which are efficiently transported to the seeds (up to ∼3% by weight^24^), potentially causing toxicity^25, 26^. By comparing the backbones of the major QAs found in NLL^14, 25^, we hypothesized that (−)-sparteine may be a common biosynthetic intermediate, and that a 2-step enzymatic oxidation would transform it into the abundant (+)-lupanine (Fig. 1a). To verify the previous reports that (+)-lupanine from NLL was enantiomerically pure^16^, we subjected plant extracts to analysis by liquid chromatography coupled to mass spectrometry (LC-MS) using a chiral column. Comparison to authentic standards confirmed that both leaves and seeds accumulated (+)-lupanine exclusively (Fig. 1b).

As genes involved in a given specialized metabolite pathway tend to be co-expressed^27^, we selected three oxidase candidates (CYP76E36, CYP71A168, and CYP71D189) whose expression patterns resembled that of the known QA pathway enzyme lysine decarboxylase (LDC) (Fig. 2a, Suppl. Table 1). We cloned the coding sequences from cDNA and expressed them individually in *Nicotiana benthamiana* via agroinfiltration. Similar to the negative control (GFP), leaves expressing CYP76E36 and CYP71A168 seemed not to be able to metabolize the separately infiltrated (−)-sparteine. By contrast, leaves expressing CYP71D189 produced a putative didehydrosparteinium ion upon infiltration with (−)-sparteine (Fig. 2b-d). We speculated that CYP71D189 was able to hydroxylate (−)-sparteine at position 2, and that, in the absence of a second, dedicated oxidase, the adjacent nitrogen eliminated the hydroxyl group spontaneously (either *in planta* or in the acidic extract^28^) (Fig. 2d).

**Figure 2.**
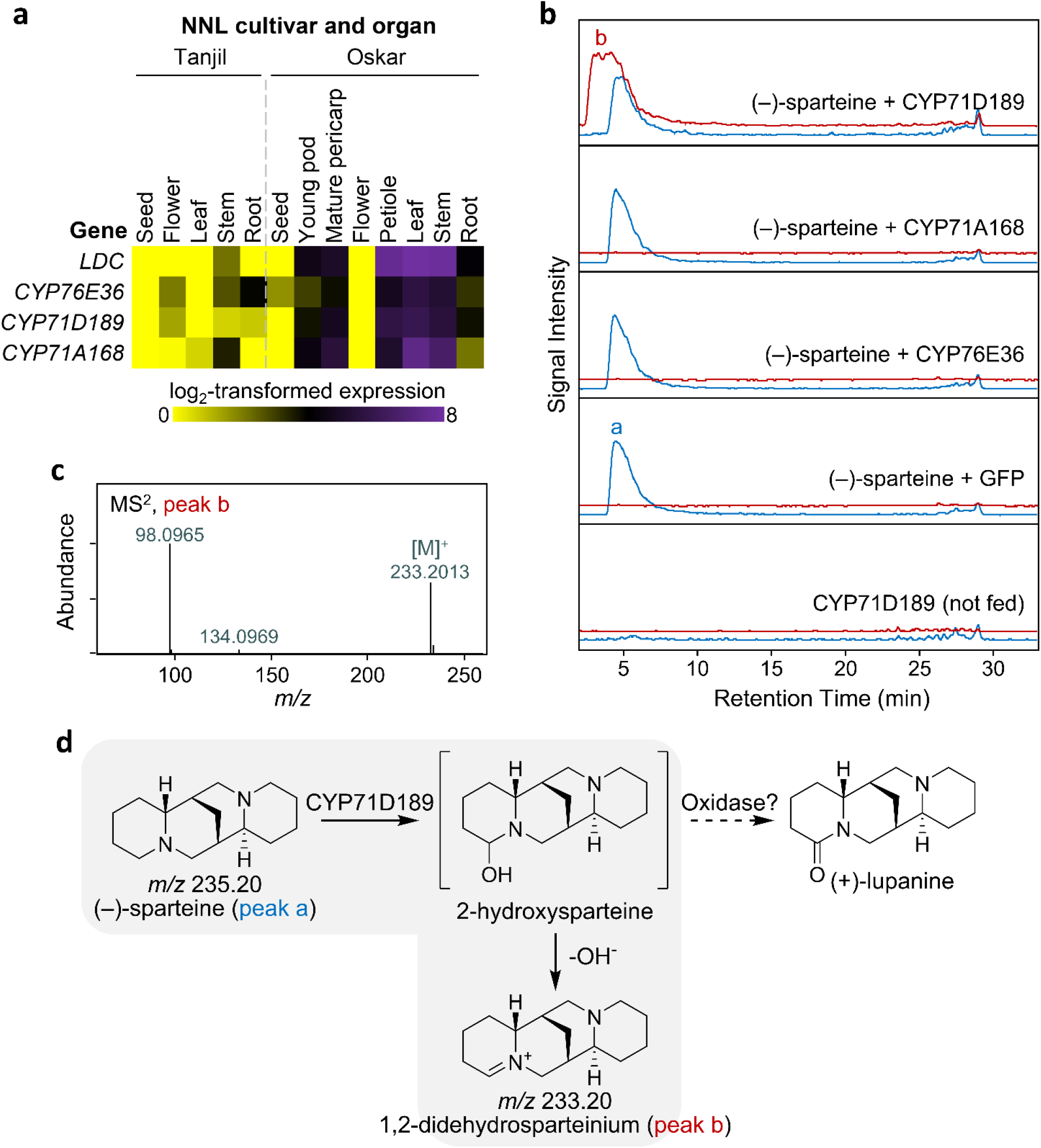
CYP71D189 is a putative sparteine 2-hydroxylase. **a**. Gene expression heatmap of the three candidate oxidases in relation to the known QA pathway enzyme LDC. Colors represent expression values in transcript per million (TPM). **b**. LC-MS analysis of extracts of *N. benthamiana* leaves expressing CYP71D189 or two other oxidase candidates at 9 days post infiltration (dpi) following feeding with (–)-sparteine at 4 dpi. Peak a corresponds to (–)-sparteine, and peak b corresponds to a didehydrosparteinium ion. Unfed leaves expressing CYP71D189 and fed leaves expressing GFP were included as controls. The traces are representative extracted ion chromatograms corresponding to sparteine ([M+H]^+^, *m/z* 235.22 ± 0.01, blue trace) and 1,2-didehydrosparteinium (M^+^, *m/z* 233.20 ± 0.01, red trace). The traces are slightly offset to aid visualization of the otherwise overlapping peaks. **b**. ESI+ CID MS^2^ mass spectrum (24.0 eV) of putative 1,2-didehydrosparteinium from CYP71D189-expressing *N. benthamiana* leaves fed with (–)-sparteine. **c**. Proposal for the CYP71D189-catalyzed hydroxylation in the absence of a subsequent enzyme-catalyzed dehydrogenation (grey background).

To find the second oxidase, we first tested CYP76E36 and CYP71A168 in *N. benthamiana* by co-expressing them individually with CYP71D189 and then feeding (−)-sparteine. Co-expression of neither cytochrome P450 changed the metabolite profile compared to a negative GFP control (Suppl. Fig. 1). Thus, we modified our selection strategy to find additional candidate genes. We focused on short-chain dehydrogenase/reductases (SDRs) given that several SDRs have been shown to aid cytochrome P450s in the oxidation of fully reduced carbons to carbonyl compounds in plant specialized metabolism^29, 30^. When applying the following three criteria, only one SDR candidate was left: 1) high expression in leaves (highly active biosynthetic organs), 2) higher expression in leaves of a high-QA variety compared to a low-QA variety, 3) absence of closely related homologs in non-QA containing plants. We named the SDR candidate SDR1 (see expression pattern in Suppl. Table 1). Gratifyingly, co-expression of SDR1 and CYP71D189 in *N. benthamiana* leaves led to decreased levels of the didehydrosparteinium ion as well as the appearance of (+)-lupanine upon infiltration with (−)-sparteine (Fig. 3a-b). We propose that SDR1 acted on the immediate product of CYP71D189, 2-hydroxysparteine (Fig. 3c).

**Figure 3.**
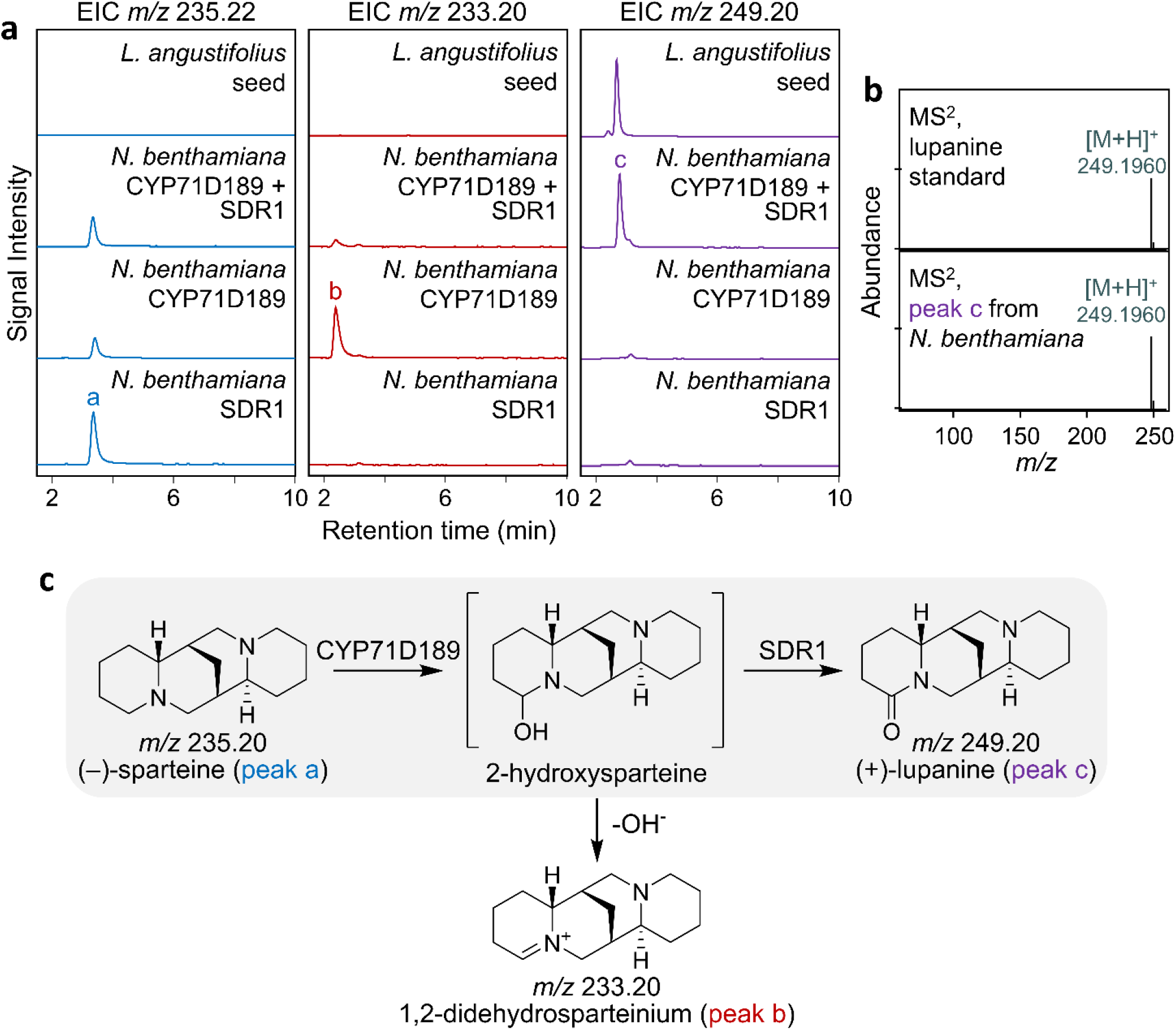
Co-expression of CYP71D189 and SDR1 enables production of (+)-lupanine from (–)-sparteine in *N. benthamiana*. **a**. LC-MS analysis of extracts of *N. benthamiana* leaves expressing CYP71D189 and SDR1 at 8 days post infiltration (dpi) following feeding with (–)-sparteine at 4 dpi. The traces are representative extracted ion chromatograms corresponding to sparteine ([M+H]^+^, *m/z* 235.22 ± 0.01, left column), 1,2-didehydrosparteinium (M^+^, *m/z* 233.20 ± 0.01, middle column), and lupanine ([M+H]^+^, *m/z* 249.20 ± 0.01, right column). Respective traces from the analysis of NLL seed extracts are also shown for comparison. Peak labels correspond to the compounds shown in panel c. **b**. ESI+ CID MS^2^ (24.7 eV) mass spectrum of lupanine produced in *N. benthamiana* leaves in comparison to a commercial standard. **c**. Proposed pathway for the stepwise oxidation of (–)-sparteine, first to 2-hydroxysparteine by CYP71D189 and then to (+)-lupanine by SDR1 (grey background).

Motivated by the prospect of generating a (−)-sparteine-accumulating NLL plant, we isolated a CYP71D189 knockout from our recently constructed, non-GMO NLL mutant library^31^ (Suppl. Fig. 2). The basis for the library was the bitter cultivar Oskar, whose seeds accumulate high levels of (+)-lupanine, (+)-13-hydroxylupanine, and (−)-angustifoline. The homozygous knockout mutants (CYP71D189^KO^) presented much reduced amounts of these major QAs in seed extracts, with only 0.6%, 2.3%, and 1.4% left, respectively. In their place, knockout seeds accumulated large amounts of (−)-sparteine, which could not be detected in extracts of wild-type seeds (Fig. 4a-b). The levels of (−)-sparteine in the extracts accounted for 1.2% of the weight of the mature seeds. For completeness, we also analyzed five minor QAs also present in Oskar seeds: (−)-multiflorine, (+)-α-isolupanine, (+)-13-hydroxy-α-isolupanine, (−)-α-isoangustifoline, and an ester of (+)-13-hydroxy-α-isolupanine. With the exception of (−)-multiflorine, all the minor QAs were strongly decreased in mutant seeds, with residual amounts ranging from undetectable [(−)-α-isoangustifoline)] to 2.4% [(+)-α-isolupanine] (Fig. 4a, c). (−)-multiflorine, however, increased 13X starting from a trace amount in wild-type seeds (Fig. 4c). In addition, we detected the appearance of a minor amount of (−)-α-isosparteine in mutant seeds (Fig. 4a, c). Based on the overall results described above, we conclude that CYP71D189 and SDR1 act in the major QA route represented by (−)-sparteine as well as in the minor one represented by (−)-α-isosparteine (Figs. 4d). Furthermore, we suggest that (−)-multiflorine is produced from either (−)-sparteine or the respective di-iminium cation precursor (Fig. 4d).Our results confirm the intermediacy of (−)-sparteine in relation to (+)-lupanine and its derivatives, which has not been clear in recent depictions of the general QA pathway^32, 33^.

**Figure 4.**
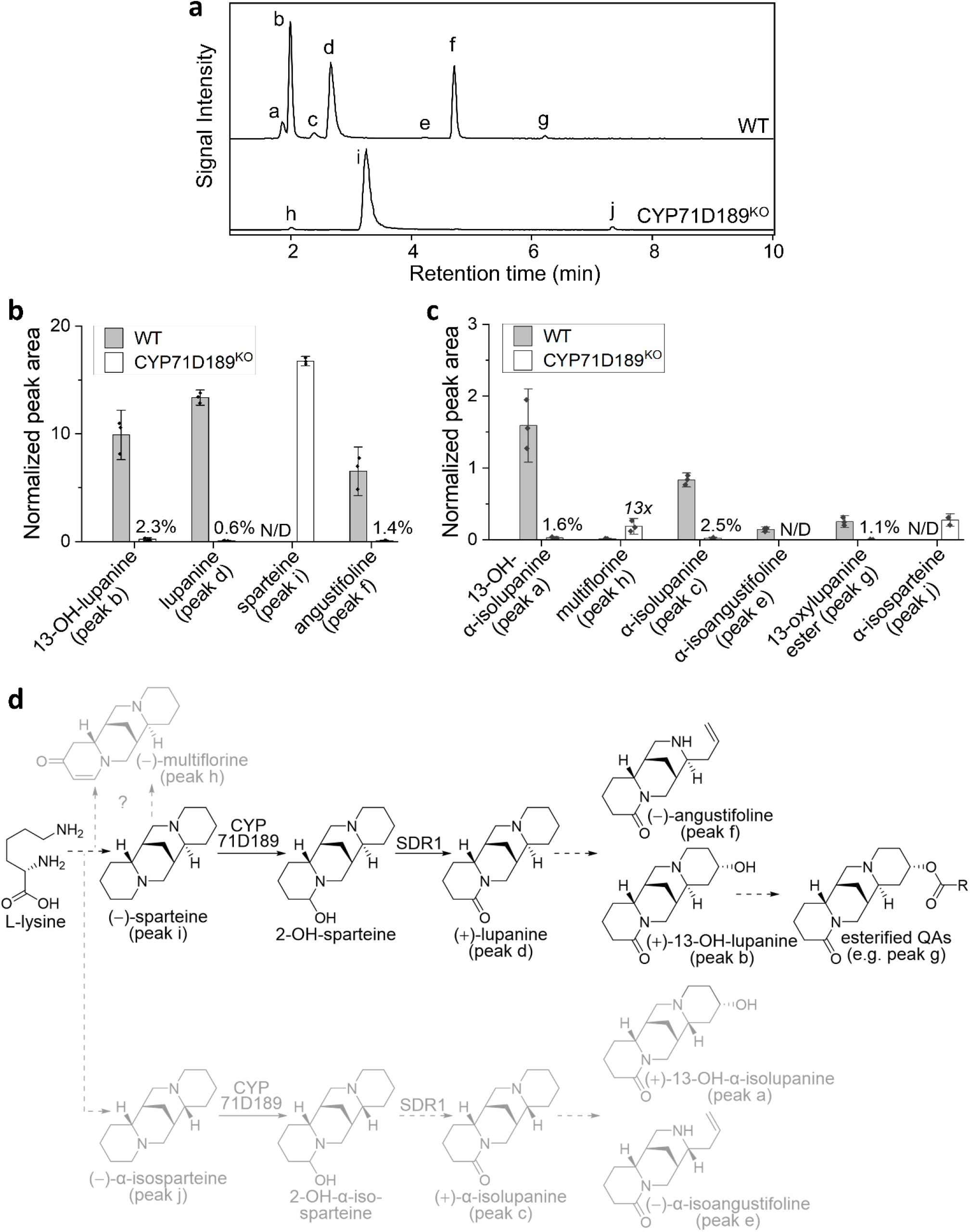
Production of (–)-sparteine in NLL by genetic inactivation of CYP71D189. **a**. LC-MS analysis of extracts of wild-type (WT) or CYP71D189^KO^ NLL seeds. Traces are representative extracted ion chromatograms at the combined *m/z* ratios of 235.18 ± 0.01 (angustifoline), 235.22 ± 0.01 (sparteine), 247.18 ± 0.01 (multiflorine), 249.20 ± 0.01 (lupanine), 265.19 ± 0.01 (13-hydroxylupanine), and 429.24 ± 0.01 (13-[3-OH-3-(4-OH-phenyl)propanoyl]-oxylupanine) ([M+H]^+^ in all cases). Peak labels correspond to compounds shown in panel d and quantified in panels b and c. **b**. Distribution and relative abundance of the four major QAs detected in WT vs. CYP71D189^KO^ seeds (*n* = 3). Data points are peak areas normalized by internal standard (caffeine) and dry seed weight. Bar charts represent mean values ± 1.5 SD. Percentages represent the residual amount of individual QAs in CYP71D189^KO^ vs. WT seeds. **c**. Distribution and relative abundance of six minor QAs detected in WT vs. CYP71D189^KO^ seeds (*n* = 3) in the style of panel b. As multiflorine was more abundant in CYP71D189^KO^ seeds, the difference compared to WT is shown as a fold change. **d**. Proposed QA biosynthesis pathway in NLL. The major QA branch is drawn in black and the minor branches are drawn in grey. Full arrows indicate steps catalyzed by known enzymes; dashed arrows indicate putative steps.

We then isolated (−)-sparteine from the relatively small M_5_ seed batch (Suppl. Fig. 2) using acid-base extraction. Compared to a crude methanolic extract, this simple procedure enabled an enrichment from 61% of total LC-MS peak area to 96% (Suppl. Fig. 3a), albeit at a yield reduction from 0.8% to 0.2% (dry seed weight). In addition, we subjected the purified (−)-sparteine to chiral GC-MS analysis, which revealed an *ee* higher than that of our commercial standard (99% compared to 97%) (Suppl. Fig. 3b). Finally, we isolated (−)-sparteine from the larger M_6_ seed batch (Suppl. Fig. 2) including crystallization as bisulfate salt. In this case, the average yield was 0.3% (dry seed weight, assuming 100% purity of the isolated salt), with a purity of 98% as analyzed by non-chiral GC-MS (Fig. 5a) and an *ee* of >99% as analyzed by chiral GC-MS (Fig. 5b).

**Figure 5.**
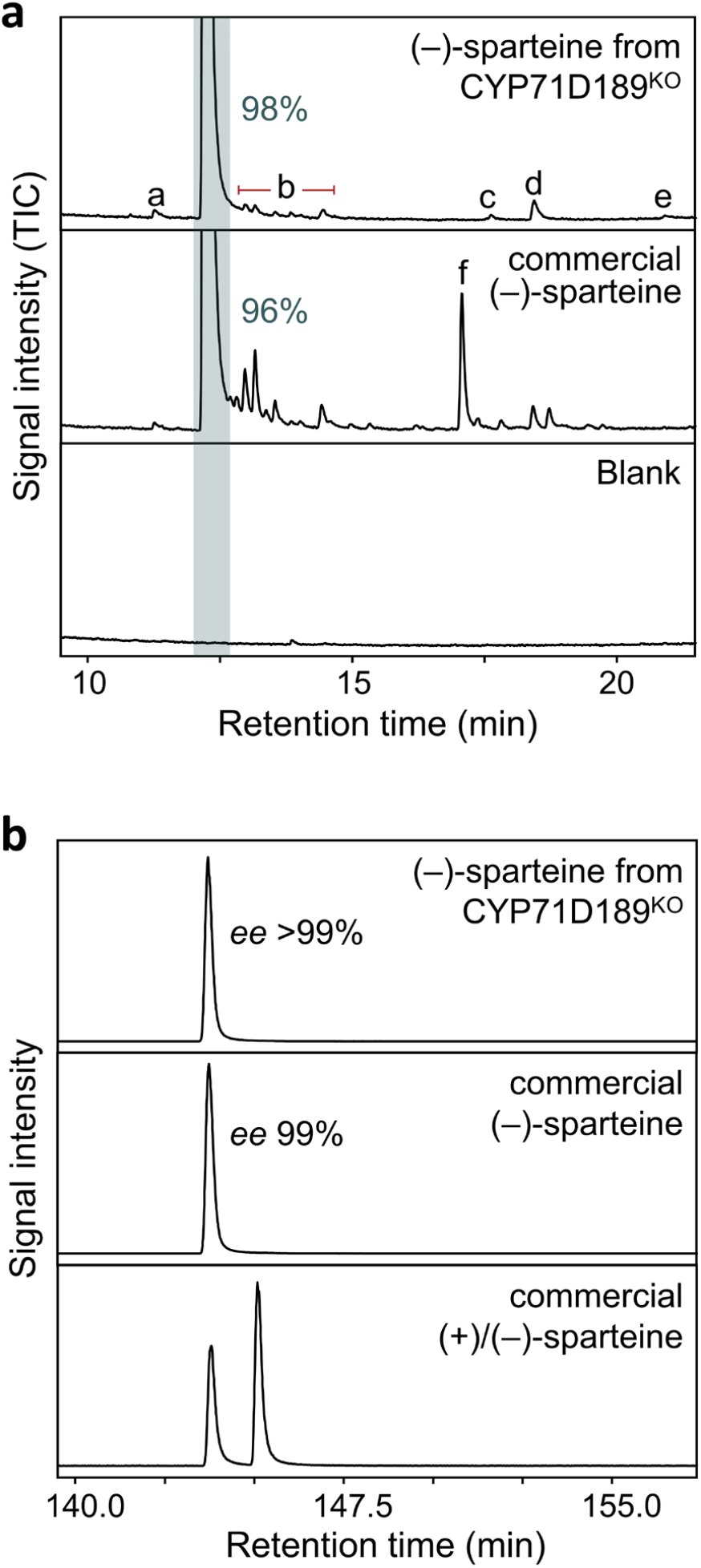
Comparison between commercial (–)-sparteine and (–)-sparteine isolated from CYP71D189^KO^ seeds. **a**. GC-MS analysis. The main contaminants of the isolated (–)-sparteine (98% pure) were α-isosparteine (peak a), various didehydrosparteine species (peaks b), α-isolupanine (peak c), lupanine (peak d), and multiflorine (peak e). In the commercial (–)-sparteine (96% pure), the main contaminant was an unknown oxosparteine species (peak f), which could have arisen from air oxidation due to prolonged storage. GC-MS traces are total ion chromatograms (TICs) and are zoomed around the baseline to show minor peaks. The sparteine peak is highlighted by the grey band. **b**. Chiral GC-MS analysis. The enantiomeric excess (*ee*) of (–)-sparteine from CYP71D10^KO^ is comparable to or higher than that of the commercial product. Chromatograms are representative extracted ion chromatograms at *m/z* 137 (base peak of sparteine).

Since our NLL mutant library was created using EMS mutagenesis, CYP71D189^KO^ plants are exempt from GMO regulations worldwide and do not require regulatory approval to be grown in the field. In our small-scale experiments, we did not observe any visible phenotypes or apparent yield penalties compared to wild-type plants. Furthermore, mature (dry) seeds can be ideal sites for long-term storage, as they may offer an extra level of protection against air oxidation. We conclude that our NLL CYP71D10^KO^ plants are a valuable source of (−)-sparteine for academia and industry.

## Notes

The authors declare being the inventors of a patent application filed by the University of Copenhagen on the subject matter of this article (PCT/EP2023/087642).

## Acknowledgements

We thank the VILLUM Foundation (project 15476) and the Novo Nordisk Foundation (projects NNF2019OC53580 and NNF22OC0075193) for their generous support.

