## Supporting Information for "Metabolic engineering of narrow-leafed lupin for the production of enantiomerically pure (‒)-sparteine"

### Contents

Supplementary Figures (1 to 3)

Gene Sequences

Materials and Methods

References

**Other Supporting Information for this manuscript**

Supplementary Tables (1 and 2)

### Supplementary Figures

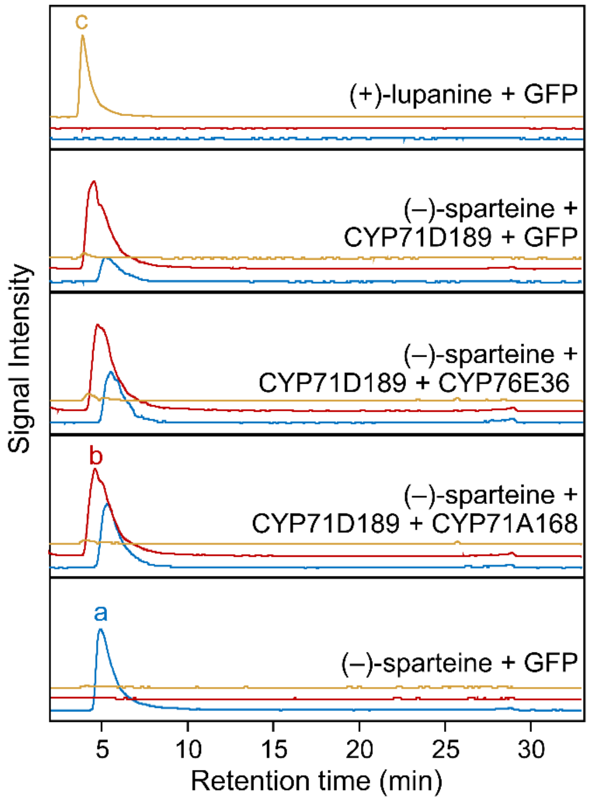


**Supplementary Figure 1 | CYP71A168 and CYP71E36 cannot oxidize 2-hydroxysparteine to lupanine.** LC-MS analysis of extracts of *N. benthamiana* leaves expressing CYP71D189 in separate combination with two other oxidase candidates at 10 dpi following feeding with (–)-sparteine at 5 days post infiltration. Peak a corresponds to (+)-lupanine, peak b to 1,2-didehydrosparteinium, and peak c to (–)-sparteine. None of the two oxidase candidates is capable of oxidizing sparteine further to lupanine. Leaves expressing GFP fed with (–)-sparteine or (+)-lupanine were included as controls. The traces are representative extracted ion chromatograms corresponding to sparteine ($[M+H]^+$, m/z 235.22 ± 0.01, blue trace), 1,2-didehydrosparteinium ($M^+$, m/z 233.20 ± 0.01, red trace), and lupanine ($[M+H]^+$, m/z 249.20 ± 0.01, orange trace). The traces are slightly offset to aid visualization of the otherwise overlapping peaks.

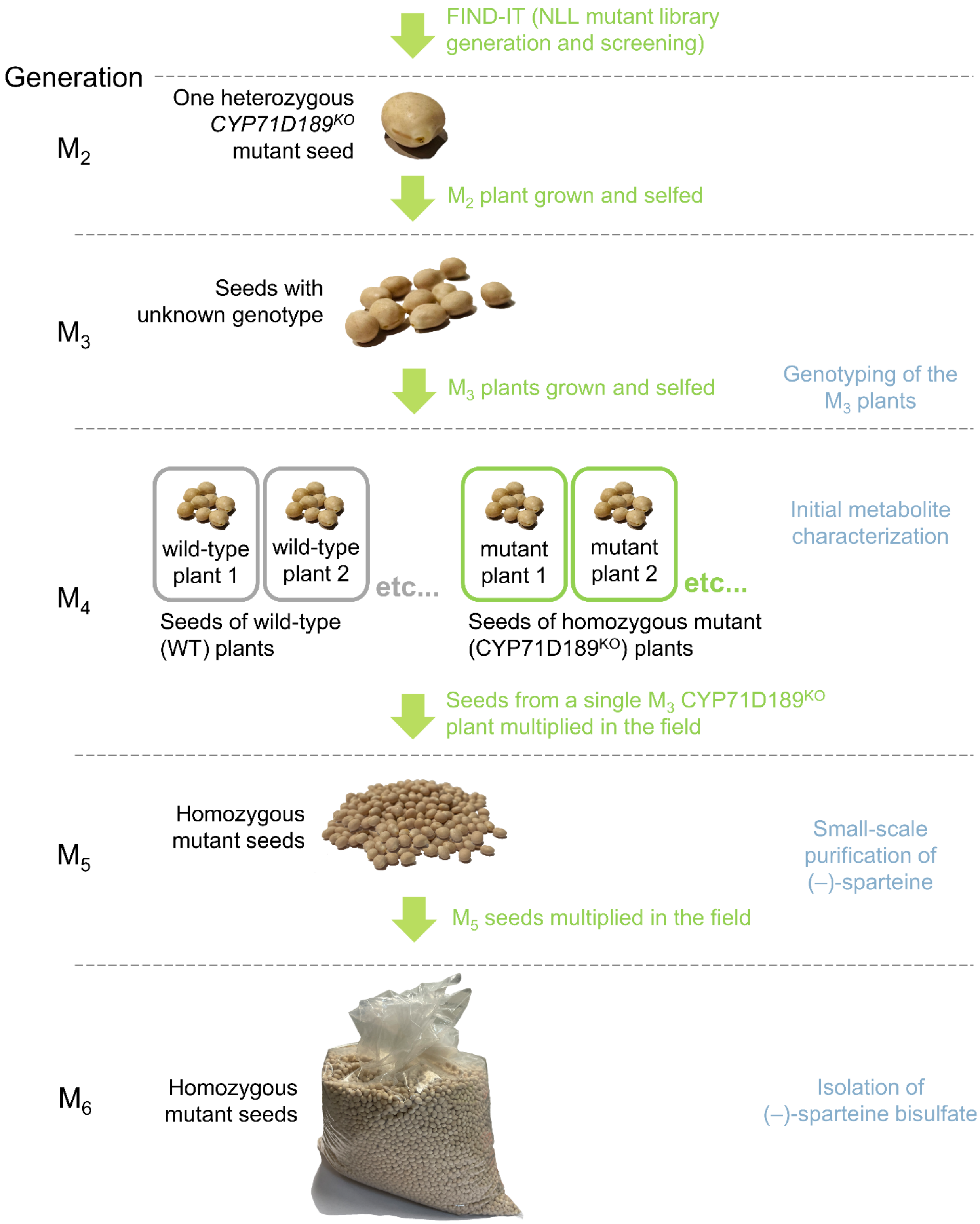


**Supplementary Figure 2 | Propagation of the CYP71D189$^{KO}$ mutant.** We retrieved one $M_2$ heterozygous *CYP71D189$^{KO}$* mutant seed from our previously constructed narrow-leafed lupin (NLL) mutant library[1] We allowed the heterozygous $M_2$ plant that grew from this seed to self-pollinate and grew a portion of the $M_3$ seeds obtained from this plant under controlled conditions. We genotyped the resulting $M_3$ plants (a mixture of wild type, heterozygous, and homozygous plants) and allowed wild-type plants and homozygous mutants (CYP71D189$^{KO}$) to self-pollinate. We then grew the $M_4$ seeds from a single $M_3$ CYP71D189$^{KO}$ plant in the field to give $M_5$ seeds, and these were further multiplied to give the $M_6$ generation.

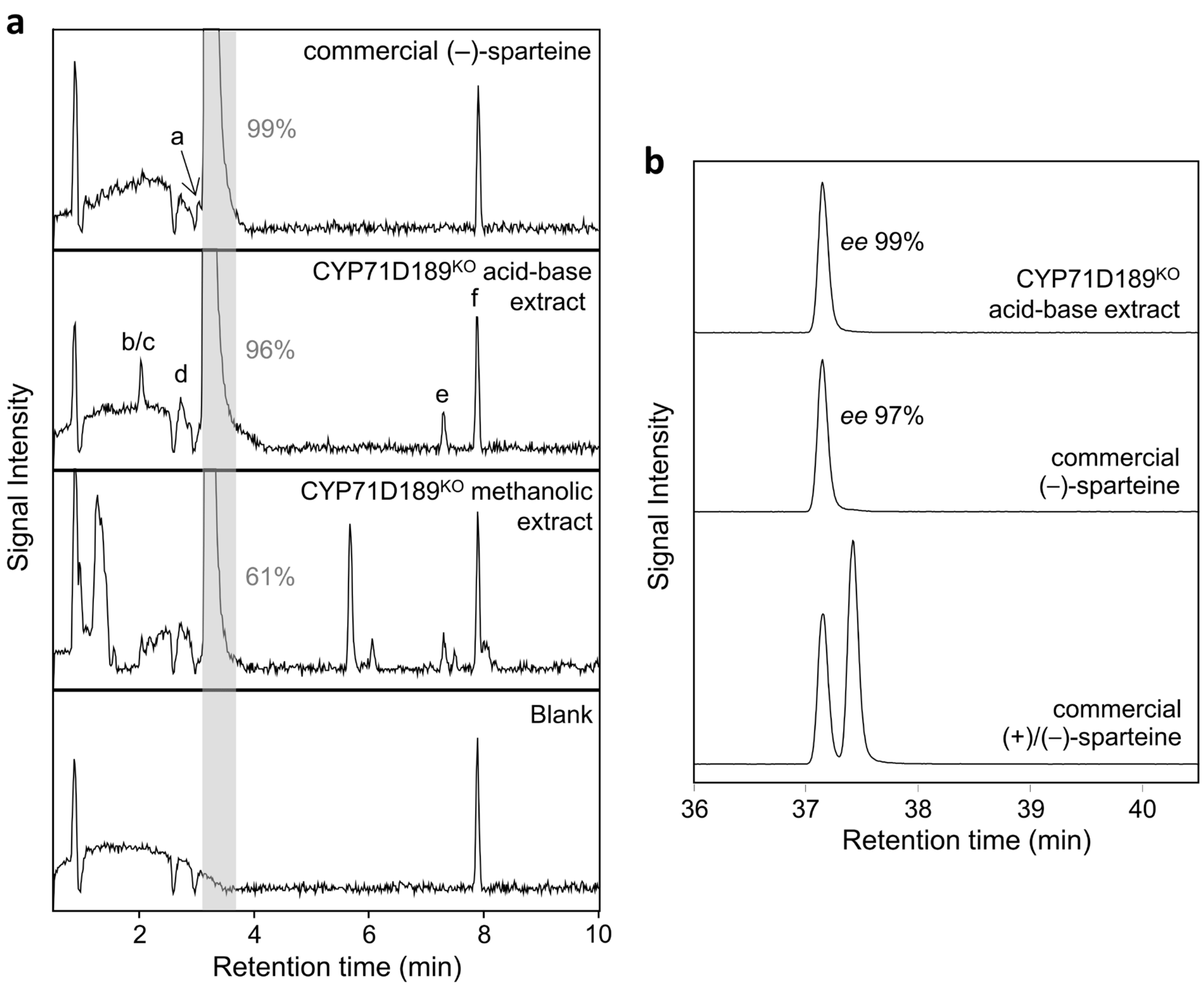


**Supplementary Figure 3 | Small-scale purification of (–)-sparteine from the $M_5$ generation of CYP71D189$^{KO}$ mutant seeds. a.** LC-MS analysis of crude methanolic extracts and extracts purified by acid-base extraction. The acid-base extraction removed non-alkaloidal contaminants, leaving mostly sparteine (96% by LC-MS peak area) and a small amount (4%) of 13-hydroxylupanine (peak b), multiflorine (peak c), lupanine (peak d) and α-isosparteine (peak e) as main contaminants. In the commercial (–)-sparteine (99% pure by LC-MS peak area), the main contaminant was found to be an unknown oxosparteine species (peak a), which could have arisen from air oxidation upon prolonged storage. LC-MS traces are total ion chromatograms (TICs) and are zoomed around the baseline to show minor peaks. The sparteine peak is highlighted by the grey band. The peak around 8 min corresponds to the internal standard (caffeine, peak e). **b.** Chiral GC-MS analysis of acid-base extracts from the seeds of CYP71D189$^{KO}$ plants compared to commercial products. The enantiomeric excess (*ee*) of (–)-sparteine from CYP71D189$^{KO}$ is comparable or higher to that of the commercial product (99% compared to 97%). Chromatograms are representative extracted ion chromatograms at m/z 137 (base peak of sparteine).

#### Gene sequences

*Sequence of CYP71D189. Exons are highlighted in grey.*

ATGGAGCTTCAAAACCCTTTCACTATTTTGTTTACATCATTTCTCTTTCTATTCTTGTTACTAGAAATAGTTAAGAGATCC
AGTTCAAAGAATTCCTATACAAACTTACCACCAGGGCCATGGAAACTACCCTTCATAGGTAACATACACCAATTTGTTG
GGTCAATGCCCCATCACTCCATGAGAGATTTAGCAGCCAAGTATGGTCCTATAATGCACCTAAAACTAGGAGAAGTTT
CCAACATCATAGTTAGTTCAGCAGAAATTGCTAGCCAGATTATGAAAACACATGATGCCAATTTTTCTTACAGGCCAGA
GAGTCTTTTTGCCAAAATATTTTCTTATAATGCTTCAGACATTGAATTCTCTCAATATGGAGATTATTGGAGGCAACTAA
GAAAGATATGCACCGTCGAGTTGCTATCAGCAAAACGTGTTCAATCATTCAGGTCATAATTATCTTTATGACTTTTCTTA
ATATTATATAATTCGGATGTGAATAAATTATTAACTGATATCTTTATTATTATATCAAAATATGTATCTTCATAAGATTTG
GTATATAGTTTACTATTGTCAGAATCTAAGATCCCACGTACATATATATATATTATCTATATGCTGGCCTCATAACTTCA
AATTTTTATGTGTGTTGCTAAATATTCAGGTTCATAAGAGAAGAGGAAGTGTCAAAACTTGCTAAAATAATATGTACAA
GTGAGGGGTCCATTGTGAATCTGTCTCCTATGATTTCCTCATTGTTGCATGGGATATCAGCACGATCAGCTTTTGGTAA
AATAAATAAAAATCAGAAAATATTGATATCAGCAATTGAGGAAGGAATTTTTCTAGCAGGAGAACTATGGGTTAGTG
AATTCTATCCCTCATTAACAGTGCTTCAAAAGTTGAGTAAGACAAAGGCTAAACTTGAAAAGTTGCACATAGAGGCTG
ATAAAATATTGCAAGAGATTCTTGATGATCACAAAAATAAAAAAAGCAGTGAGAGTGAAGATCTAATAGATGTTCTTC
TCAAGTTTCAAAATGACAAGGATTCTCAACCTCCCTTAACTGATGACAATATTAAAGCAATTGGCCAGGTTAGTACATT
TTCTCTATGTATGTCTAACATTATTCTTTATGTGTAAAAATTAGGTAGAATTTATAACATTATTTATGATGAATGATTTTA
ATCAATCTCCATGAAATGATTGGTGTCCTCGATTATGTAATAAAGATTCATGTTTGTATGGAAGAAAATGTTTATGGAT
AAGGGATTATGAATCCATTTATTATAGGAGATGTTTGGTGGTGGTGGTGAAACAACATCAAGTACTGTTGTGTGGTGT
TTGTCAGAAATGATAAAGAAACCAAAAGTGATGGAAGAAGCACAAGCTGAGGTAAGAAAAGTGTATGATAATAAAG
GATATGTGGATGAGTCAGAGTTGCACCAATTGATATATTTGAAGGCAGTGATCAAAGAAACACTGAGGCTCCACCCAC
CTGTCCCATTGTTAATGCCAAGAGAAAATAAAGATAGCAGCAAAATCAATGGATATGATATACCACCTAAGTGCAAGG
TCTTAATAAATGCTTTTGCTATTGGAAGAGATCCTAAGTATTGGAATGAACCTGAAAGCTTTAATCCTGAGAGATTTCT
GAATAGTTCAGTAGATTATAAGGGCACAGACTTTGAGTTTATACCATTTGGTGCTGGAAGGAGGATATGCCCTGGTAT
TACATTTGCCATACCAAACATGGAGCTTCCACTTGCAACATTGCTTTACCATTTTGATTGGAAGCTTCCAAATGGAATG
AAGAATGAAGAACTTGATATGGATGAATCATCTGGGTTGGCTATTAAAAGAAAAAATGATCTTTGCTTGATTCCAATT
GTTACTCGTATGCCTTAA

*Sequence of CYP71D189$^{KO}$. Exons are highlighted in gray. The nonsense mutation is highlighted in blue.*

ATGGAGCTTCAAAACCCTTTCACTATTTTGTTTACATCATTTCTCTTTCTATTCTTGTTACTAGAAATAGTTAAGAGATCC
AGTTCAAAGAATTCCTATACAAACTTACCACCAGGGCCATGGAAACTACCCTTCATAGGTAACATACACCAATTTGTTG
GGTCAATGCCCCATCACTCCATGAGAGATTTAGCAGCCAAGTATGGTCCTATAATGCACCTAAAACTAGGAGAAGTTT
CCAACATCATAGTTAGTTCAGCAGAAATTGCTAGCCAGATTATGAAAACACATGATGCCAATTTTTCTTACAGGCCAGA
GAGTCTTTTTGCCAAAATATTTTCTTATAATGCTTCAGACATTGAATTCTCTCAATATGGAGATTATTGGAGGCAACTAA
GAAAGATATGCACCGTCGAGTTGCTATCAGCAAAACGTGTTCAATCATTCAGGTCATAATTATCTTTATGACTTTTCTTA
ATATTATATAATTCGGATGTGAATAAATTATTAACTGATATCTTTATTATTATATCAAAATATGTATCTTCATAAGATTTG
GTATATAGTTTACTATTGTCAGAATCTAAGATCCCACGTACATATATATATATTATCTATATGCTGGCCTCATAACTTCA
AATTTTTATGTGTGTTGCTAAATATTCAGGTTCATAAGAGAAGAGGAAGTGTCAAAACTTGCTAAAATAATATGTACAA
GTGAGGGGTCCATTGTGAATCTGTCTCCTATGATTTCCTCATTGTTGCATGGGATATCAGCACGATCAGCTTTTGGTAA
AATAAATAAAAATCAGAAAATATTGATATCAGCAATTGAGGAAGGAATTTTTCTAGCAGGAGAACTATGAGTTAGTGA
ATTCTATCCCTCATTAACAGTGCTTCAAAAGTTGAGTAAGACAAAGGCTAAACTTGAAAAGTTGCACATAGAGGCTGA
TAAAATATTGCAAGAGATTCTTGATGATCACAAAAATAAAAAAAGCAGTGAGAGTGAAGATCTAATAGATGTTCTTCT
CAAGTTTCAAAATGACAAGGATTCTCAACCTCCCTTAACTGATGACAATATTAAAGCAATTGGCCAGGTTAGTACATTT
TCTCTATGTATGTCTAACATTATTCTTTATGTGTAAAAATTAGGTAGAATTTATAACATTATTTATGATGAATGATTTTAA
TCAATCTCCATGAAATGATTGGTGTCCTCGATTATGTAATAAAGATTCATGTTTGTATGGAAGAAAATGTTTATGGATA
AGGGATTATGAATCCATTTATTATAGGAGATGTTTGGTGGTGGTGGTGAAACAACATCAAGTACTGTTGTGTGGTGTT
TGTCAGAAATGATAAAGAAACCAAAAGTGATGGAAGAAGCACAAGCTGAGGTAAGAAAAGTGTATGATAATAAAGG
ATATGTGGATGAGTCAGAGTTGCACCAATTGATATATTTGAAGGCAGTGATCAAAGAAACACTGAGGCTCCACCCACC
TGTCCCATTGTTAATGCCAAGAGAAAATAAAGATAGCAGCAAAATCAATGGATATGATATACCACCTAAGTGCAAGGT
CTTAATAAATGCTTTTGCTATTGGAAGAGATCCTAAGTATTGGAATGAACCTGAAAGCTTTAATCCTGAGAGATTTCTG
AATAGTTCAGTAGATTATAAGGGCACAGACTTTGAGTTTATACCATTTGGTGCTGGAAGGAGGATATGCCCTGGTATT
ACATTTGCCATACCAAACATGGAGCTTCCACTTGCAACATTGCTTTACCATTTTGATTGGAAGCTTCCAAATGGAATGA
AGAATGAAGAACTTGATATGGATGAATCATCTGGGTTGGCTATTAAAAGAAAAAATGATCTTTGCTTGATTCCAATTG
TTACTCGTATGCCTTAA

*Coding sequence of SDR1*

ATGGTAGAAACTACTTCCAACAACAGTGGTGCAAAGCTAGCAGGCAAAGTAGCCATCGTCACCGGAGGATCCGGTGG
CATCGGCGAGGCAGCGGCGCATGCCTTTGCTGATCAAGGTGCGCGTGTGGTAATTGCAGACGTTCAAGACGATCTTG
GCAATAAAGTTGCTGAGTCCATCGGAACCAACAAGTGTACCTACATCCACTGCAACGTAGCAGATGAAGAACAAGTCC
AAAACCTAATTCAATCAACGGTCAACACTTTCGGACAAATCGACATCATGTTCAGCAACGCTGGCATCAGCAGTGAGT
TAGAACAAACCATCATTGAACTTGACATATCCGAACTTAACCGCTTGTTGGCCGTAAACGCTAGTGGAATGGCGGCGT
GTGTGAAACACGCGGCACGTGCCATGGTGGACAAGCACGTGAGAGGAAGCATAGTATGCACTGGAAGCATATATGG
AAGTAACGGTGGATCTGATGGAACGGATTACGCCATGTCGAAGCATGCATTGTTGGGTTTGGTGCGTTCGGCGAGTA
TACAACTAGCGGAGCATGGAATAAGGGTGAACTGCGTTTCACCAAATGCATTGGTAACTCCGATGACGTTGAAATATT
CAGGGAGTGAAGAGAAGGTTTATGAGCTTTGTGCAAAAACCGCGAGGCTGAAAGGAGTGATACTCACTGCTAAACAT
ATTGCTGATGCTGTGTTATTTCTTGCTTCTAATGATTCAGAGTTTATCACTGGCCATGATCTTGTTGTGGATGGTTCCTA
TATTGCTCCATGA

*Coding sequence of CYP76E36*

ATGGATTATCTAACACTTTTTCTACTCATTTCCTTTGTTTGGACAAGCATTTATGTCCTATTCTCCAAATTAGGAATCAAA
ACATCCAAATATGCCCCAGGTCCATACCCTTTACCTATCATAGGTAACATCTTTGAACTTGGAAAACTCCCACACCAAAC
ACTTTCTAAGCTCTCTCAAACTTATGGACCTATAATGACCTTAAAGTTTGGTAGTGTTACTGCTATAGTTATTTCCTCTCC
ACAAGTAGCCAAAGAAGCACTTCAAAAAAATGACCAAGTTTTCTCTTTTAGGCCAACCCCAGATACCCTTAGGGCACA
TGACCATCATATATACTCAGTGGCATGGATGCAGCCTTCAGCTGATTGGAGGGCCCTTAGGAAAGCTTGCGCAATCAA
AGTGTTCTCATCTCGAATGCTCGATTCGACGCAATTTTTACGACAAAAGAAGGTGCAAGAGTTGATGGATTATGTTAA
GGAAAGTTGCAAGAAAGGTGAGGCTTTGGATATTGGCAAGGCAACTTTTAAGACTGTGCTTAATTCTATATCAAACAC
TTTGTTCTCTATGGACTTGGCTCATTATACTTCTGATAAGTTTCAAGAGTTCAAGGACATTATTTGTGGGATCACTGAAG
AAGCTGGAAAGCCTAACTATGTGGATTATTTTCCAATCCTTAGTTTTCTTGATCCACAAGGTGCCCATGGAAGAATGAA
GGGTTATTTTGGAAAGTTGATTAAATTTTTTGATGATCTTATAGAAGAAAGGCTACAATTAAGAGCTACACAAAAGGA
ATCCAAGGCTTGCAAAGATGTTCTAGATTCTGTGCTAGAACTCATGCTGGAAGACAATTCTCAAATTACTAGGCTCCAT
GTTTCGCATTTGTTTGTGGATTTATTCGTGGCTGGAATAGATACCACATCAATCACAATAGAATGGGCAATGGCAGAG
TTGCTACGTAATCCAGAAAAGCTAAAAAAAGTTAGAAAAGAACTTCAACAAGTTACAAGCAAAGGTGAACAACTTGA
AGAAACACACATATCAAAGCTTCCTTTCTTAGAAGCAGTGATTAAAGAAACTTTTCGTTTGCATCCACCAGCAGCATTC
TTAGTGCCACGCATGTCAGGAGATAATGTTGAACTATGTGGTTACATGGTACCTAAAAATGCACAAATTATGATTAAT
GCATGGGCCATGGGAAGAGATTCAAGTGTTTGGGCCAACCCAAATGAATTTATCCCTGAAAGATTCTTGAATAATGAG
ATTGATTTTAAAGGTCAATATTTTGAGCTTATTCCTTTTGGTGCTGGAAGAAGGATTTGTCCTGGTTTACCATTGGCTTC
TAAGACTGTGCACACTGTCTTGGCCTCACTTTTATGTGGCTATGATTGGAAGCTTGTTGATGGAGGAGAGGGAGAGA
ATATGGATATGTCTGAGGAATATGGGCTTACCTTACATAAGGCACAACCTCTCCTAGTTATTCCTATCAAAGCATAA

*Coding sequence of CYP71A168*

ATGCCTCTTCTATCTATTCAAGAGAATCTACCATATGTGCTAAATTCAACTTTCTACTTATCTGTACTATGTTTCTTTGGT
GTCTTATTTGTCTTTAAGATCACTAGAAGAAGCAAAACCAATTCACCACCCTCCCCTCCAAAACTACCCTTTATTGGGAA
TCTTCATCAACTAGGCACATTCCCACACCGTTCCTTACAATCCTTATCTTACAAATATGGCCCTATGATGATGATGAAAA
TGGGACAAATTCAAACGCTAGTGATTTCATCTTCTGATGTGGCCAGAGAAATATTCAAAAGCCATGATGCTGTTTTCTC
CAACCGACCCACGGTCACAGGCTCCGACATCTTCTTGTATGGGTCCAAAGATGTGGCCTTTGCTCCCTACGGCGATGA
GTGGAGACAAAAAAGAAAGATTTTAGTTCTTGAGCTTCTAAGCATGAAAAGGGTGCAATCTTTTCAACCCATAAGAGA
AGAAGAAGTTGGTGAAATGCTTCATGCTATACGTGATGCATGCAGAAAGTCATCAACGGTGAACCTGACTGAGATGC
TGATTGCAGCCTCTAACAACTTAAATTCTAGATGTGTTTTCGGACAAAAGTATGATACTGAAGATGGCAGTCCCAGCTT
CGGAGATCTAGGAAGGAAGATGTTGGTACAGTTCACAGCTTTCTGTGTTGGAGATTTCTGGCCTTCATTGAGTTGGAT
TGATACACTTTCAGGCCAAATTCCAAAATTTATGGAAACTTTTACTTCGTTAGATATTTTCCTAGAAAGAGTTATTAAAG
AACACAGGGCTAAGATGAAGAGTAGTGATGATCAATCCGATAAGAAAGACTTCGTGGATATACTTCTTCAACTTCAAG
GAGAAGATAAGCTTGACTTTGAGCTCACCCAAGATATCCTCAAAGCACTAATAGTGAACTTGTTCATTGGAGGAAGTG
ATACTTCATCAACAACAATGGAATGGGCTTTTGCAGAACTCATGAGGAATCCAAGGGTCCTGAAGAAAGCCCAAGAA
GAGGTAAGAAGAGTTGTGGGGGACAAAAAAGTTGTAGATGCAAATGATACAAAACATATGAATTACTTGAAATGTGT
AATCAAAGAAACTTTAAGATTACATCCACCAGCTCCTCTCTTGGTTCCTAGAGAAACAACTGCTACTGTTAATCTAAAA
GGATATGACATTCCCTCCAAAACAAGGATACTTATAAATGGATTTGCAATCCAAAGGGACCCTGAAGTTTGGGACAAA
GCTGATGAGTTTTACCCAGATAGATTCGAGAACAGTGAGGTTGACTTCAAAACACAAGACGTAGAATTTATAGCATTT
GGCGGTGGAAGAAGGGGGTGCCCTGCAATTACATTTGCTGTTACCTTTACTACTTATGTGCTTGCTAATCTTCTATATT
GGTTTGATTGGAAGCTTCCTGAAAACGTAGACGAGGTGGACATGAGTGAGAGATATGGAATTGTTGTCAACTTGAAA
GTACCACTTCAACTGAAACCGGTGCTATCCTCCTTTGGAAGTGGATCTCAGCCTTGA

### Materials and Methods

*Analytical methods*

*LC-MS.* LC-MS analyses were carried out on a Thermo Fisher Dionex 3000 RS HPLC/UPLC system interfaced to a Bruker compact QqTOF mass spectrometer through an ESI source. Two different LC methods were used as described below. ESI mass spectra (*m/z* 50-1000) were acquired in positive ionization mode with automatic $MS^2$ acquisition using the following parameters: capillary voltage 4500 V; end plate offset −500 V; source temperature 250 °C; desolvation gas flow 8.0 L/min; and nebulizer pressure 2.5 bar. $N_2$ was used as desolvation, nebulizer and collision cell gas. Sparteine and lupanine were identified by comparison with known standards. (+)- and (–)-sparteine were purchased from Sigma-Aldrich. (+)- and *rac*-lupanine were purchased from Innosil (Poznan, Poland). The identity of the other QAs was inferred from their predicted molecular formula and their mass spectral pattern as shown by Otterbach *et al.*[2].

*LC method 1* (screen of oxidase candidates in *N. benthamiana* fed with (–)-sparteine). Analytes were separated at 40 °C on a Kinetex XB-C18 column (100 x 2.1 mm, 1.7 µm, 100 Å, Phenomenex). Mobile phases A and B consisted of 0.05% formic acid in water and 0.05% formic acid in acetonitrile, respectively. Analytes were eluted using the following gradient at a constant flow rate of 0.3 mL/min: 0–1 min, 2% B (constant); 1–16 min, 2–25% B (linear); 16–24 min, 25–65% B (linear), 24–26 min, 65–100% B (linear); 26–27 min, 100% B (constant); 27–27.5 min, 100–2 % B (linear); and 27.5–33 min, 2% B (constant).

*LC method 2* (test of CYP71D189 with or without SDR1 in *N. benthamiana* fed with (+)- or (–)-sparteine; QA quantification in seeds of the $CYP71D189^{KO}$ NLL mutant). Analytes were separated at 40 °C on a Luna C18(2) column (150 × 2 mm, 3 µm, 100 Å, Phenomenex). Mobile phases A and B consisted of, respectively, 0.05% formic acid in water and 0.05% formic acid in acetonitrile. Analytes were eluted using the following gradient at a constant flow rate of 0.3 mL/min: 0–0.5 min, 2% B (constant); 0.5–2.375 min, 2–6% B (linear); 2.375–7 min, 6–25% B (linear), 7–13 min, 25–100% B (linear); 13–14 min, 100% B (constant); 14–14.5 min, 100–2% B (linear); and 14.5–20 min, 2% B (constant).

*GC-MS.* Non-chiral GC-MS analysis was carried out on a Shimadzu GCMS-QP2010 Plus single quadrupole gas chromatograph-mass spectrometer equipped with a Shimadzu AOC-5000 autosampler. Analytes were separated on an Agilent J&W HP-5ms Ultra Inert capillary column (30 m x 0.25 mm, 0.25 µm) using He as carrier gas. Ethyl acetate extracts were injected in splitless mode (1 µl) at an inlet temperature of 250 °C. Analytes were separated using the following column temperature program at a constant carrier gas linear velocity of 33.7 cm/s: initial 80 °C, hold for 3 min; ramp to 150 °C at 30 °C/min; ramp to 300 °C at 6 °C/min; hold for 10 min. The separated analytes were ionized using an electron impact ion source at 250 °C. MS spectra were acquired in scan mode (m/z 30-600) at an energy of 70 eV. The purity of the extracted sparteine was estimated using the %area normalization method. In the calculation of the total peak area, we included all chromatographic peaks visible in the total ion chromatogram of the extract but absent in that of a blank.

*Chiral GC-MS analysis.* Chiral GC-MS was carried out on a Shimadzu Nexis GC-2030 gas chromatograph equipped with a Shimadzu AOC-6000 autosampler and coupled to a Shimadzu GCMS-QP2020 NX single quadrupole mass spectrometer. Analytes were separated on an Agilent J&W CycloSil-B capillary column (30 m x 0.25 mm x 0.25 µm) using He as carrier gas. Ethyl acetate extracts were injected in splitless mode (1 µl) at an inlet temperature of 250 °C. For the chiral analysis of (–)-sparteine extracted from the $M_5$ generation of CYP71D189$^{KO}$ seeds, analytes were separated using the following column temperature program at a constant carrier gas pressure of 51.0 kPa: initial 80 °C; hold for 3 min; ramp to 125 °C at 30 °C/min; ramp to 240 °C at 2 °C/min. For the chiral analysis of (–)-sparteine bisulfate isolated from the $M_6$ generation of CYP71D189$^{KO}$ seeds, analytes were separated using the following column temperature program at a constant carrier gas linear velocity of 33.7 cm/s: initial 80 °C; hold for 3 min; ramp to 125 °C at 30 °C/min; hold for 120 min; ramp to 240 °C at 2 °C/min. The separated analytes were ionized using an electron impact ion source at 250 °C. MS spectra were acquired in scan mode (m/z 10-300) at an energy of 70 eV. (+)- and (–)-sparteine were identified by comparison with commercial standards (Sigma-Aldrich). The enantiomeric excess of (–)-sparteine was calculated from the extracted ion chromatogram of *m/z* 137 (base peak of sparteine).

*Selection of gene candidates*

For gene candidate selection, we used an NLL transcriptomics dataset including eight different organs of the bitter cultivar Oskar (NCBI BioProject PRJNA386115) and five different organs of the sweet cultivar Tanjil (NCBI BioProject PRJNA248164). This includes biosynthetic as well as non-biosynthetic organs for both cultivars. The RNA-Seq reads from both cultivars were mapped onto the Oskar transcriptome[3] using Kallisto[4] and gene expression was quantified in transcripts per million (TPM).

The selection of candidate genes for the oxidation of sparteine was done by co-expression analysis using *LDC* as bait. We first reduced our dataset by removing all those transcripts with low expression (<10 TPM) in young Oskar leaves. Then, we calculated Pearson's correlation coefficients (PCCs) between *LDC* and all other transcripts. The transcripts with PCC higher than 0.9 were subjected to BLASTX against green plant protein sequences from NCBI (taxid: 33090). We distinguished between general and specific transcripts based on the %ID of the first 30 BLASTX results. Specifically, we only retained specific transcripts defined as those with 10 or more BLASTX results with %ID<70%. We searched among the annotations of the remaining transcripts for keywords indicating oxidative enzymes, thus narrowing down to three hits. The three transcripts were predicted to encode cytochrome P450s, specifically a *CYP71D*, a *CYP76E*, and a *CYP71A* (Supplementary Table 1).

To obtain additional candidates for the oxidation of 2-hydroxysparteine, we expanded our scope to include genes that were less strongly co-expressed with LDC yet had higher expression in Oskar vs. Tanjil. Hence, we filtered our dataset by removing all those transcripts with low expression (<10 TPM) in young Oskar leaves and with higher expression in leaves of Tanjil vs. Oskar (TPM ratio>1). The remaining transcripts were subjected to BLASTX and selection of specific transcripts based on %ID as described above. Finally, we searched for the keyword "dehydrogenase" among the annotations of the remaining transcripts and obtained only two hits. One of them appeared to encode a fragment of subunit CRR3 of the NAD(P)H dehydrogenase complex and was therefore deemed unlikely to be involved in the biosynthesis of QAs. The other transcript was predicted to encode an SDR specifically annotated as (–)-isopiperitenol/(–)-carveol dehydrogenase (Supplementary Table 1).

*Transient expression of biosynthetic gene candidates in* Nicotiana benthamiana

Total RNA was extracted from young leaves of narrow-leafed lupin (NLL) cv. Oskar using the Spectrum Plant Total RNA Kit (Sigma-Aldrich) and cDNA was synthesized using the iScript cDNA Synthesis Kit (Bio-Rad). Full-length coding sequences of *CYP71A168*, *CYP71D189*, *CYP71E36*, and *SDR1* (see *Gene sequences* section) were amplified from the NLL leaf cDNA using the primers shown in Supplementary Table 2. The coding sequence of mGFP5 (GFP) was amplified from the plasmid pCAMBIA1302 using the primers shown in Supplementary Table 2. The genes were cloned into the plant expression vector pEAQ-USER[5] by USER cloning[6] and transformed into *E. coli* strain Top10. Positive clones were verified by Sanger sequencing and transformed into *Agrobacterium tumefaciens* strain AGL-1. For agroinfiltration, *Agrobacterium* strains were grown in YEP liquid medium supplemented with 50 µg/mL kanamycin, 25 µg/mL rifampicin and 50 µg/mL carbenicillin at 28 °C and 220 rpm until $OD_{600} \approx 3\text{-}4$ in conical flasks. Bacterial pellets were harvested by centrifugation and resuspended in ultrapure water to $OD_{600} = 1$. Different strains were mixed in equal portions to obtain the desired combination of genes for co-expression in the leaves of *N. benthamiana*. A strain expressing GFP was used as negative control. After 1-3 h of incubation at room temperature, the mixtures were infiltrated into the abaxial side of young leaves of 4-week-old *N. benthamiana* plants using a needle-less 3-ml syringe. Infiltrated plants were allowed to recover overnight in the dark before being taken back into the greenhouse. To feed (–)-sparteine, (+)-sparteine, and (+)-lupanine, a 50 ppm solution was prepared in buffer (10 mM $MgCl_2$, 10 mM Na-MES buffer pH 5.6), and the solution was infiltrated into the abaxial side of previously agroinfiltrated leaves at 4 or 5 days post agroinfiltration. Care was taken to infuse the solutions into the entirety of the previously agroinfiltrated area, which appeared discolored. The fed plants were grown for an additional 4 or 5 days. Two leaf discs of 1-cm diameter were harvested from the agroinfiltrated (discolored) portions of each leaf at 8 to 10 days post agroinfiltration. The leaf discs were either quickly frozen in liquid nitrogen (screening of the three CYP candidates in leaves fed with (–)-sparteine) or dried overnight in an oven at 40°C (all other experiments). The frozen or dried leaf discs were pulverized using steel beads and a

TissueLyzer bead beater (Qiagen). After removal of the beads, the leaf powder was extracted with 250 µl of extractant (60% methanol, 0.06% formic acid, and 5 ppm caffeine in water) for 3 hours at 1200 rpm. The extracts were briefly centrifuged to remove leaf debris, diluted 5x with ultrapure water, and filtered through a 0.22-µm filter. The filtered extracts were analyzed by LC-MS as described above under *Analytical methods*.

*Isolation and initial characterization of CYP71D189$^{KO}$ NLL plants*

The NLL mutant library constructed previously[1] was screened essentially as described in Knudsen *et al.* (2022)[7] to identify a mutant seed harboring the specific G to A nucleotide change at position 648 in the coding region of *CYP71D189* (corresponding to W216Stop; see the *Gene sequences* section above). For the TaqMan assays, we used the following primers: a target-specific forward primer (CYP71D189_TaqMan_FW), a target-specific reverse primer (CYP71D189_TaqMan_RV), a WT-specific probe containing a HEX fluorophore and a BHQ1 quencher (CYP71D189WT_TaqMan_HEX), and a mutant specific probe containing a FAM fluorophore and a BHQ1 quencher (CYP71D189KO_TaqMan_FAM) (Supplementary Table 2). One heterozygous mutant seed was retrieved. The $M_2$ heterozygous plant that grew from the seed was allowed to self-pollinate. The resulting $M_3$ seeds were sown in 16 cm-wide, 20 cm-deep pots filled with commercial peat-based potting soil and grown in a growth cabinet with a light/dark photoperiod of 16/8 h, a day/night temperature of 21/18 °C, and a relative humidity of 60%. Young leaves from the $M_3$ plants were quickly frozen in liquid nitrogen and pulverized with the help of steel beads and a TissueLyzer bead beater (Qiagen). Genomic DNA was extracted from the leaves using the E.Z.N.A.® Plant DNA DS Kit (Omega Bio-tek), and genotyping was carried out by Sanger sequencing of a PCR fragment spanning the entire gene between its start and stop codons using primers CYP71D189_pEAQ_FW and CYP71D189_pEAQ_RV (Supplementary Table 2). All the $M_3$ WT plants and homozygous CYP71D189$^{KO}$ plants were allowed to self-pollinate. For QA analysis, three mature, dry $M_4$ seeds from either three WT plants or three homozygous CYP71D189$^{KO}$ plants were pulverized using a steel ball with the help of a TissueLyzer bead beater (Qiagen). QAs were extracted from ~20 mg of seed flour in 1 ml of extractant (60% methanol, 0.06% formic acid, and 15 ppm caffeine in water) for 3 hours at 1200 rpm. The extracts were briefly centrifuged to remove solid residues, diluted 15x with ultrapure water, and filtered through a 0.22-µm filter. The filtered extracts were analyzed by LC-MS as described above under *Analytical methods*.

*Purification and chiral analysis of (–)-sparteine from $M_5$ CYP71D189$^{KO}$ seeds*

Fifty $M_4$ seeds from one $M_3$ homozygous CYP71D189$^{KO}$ plant were grown in the field at Nørre Aaby, Fyn, Denmark. The seeds were hand sown in two separate rows 25 cm apart during April of 2022. 44 plants reached maturity and were harvested in late August 2022, yielding 438 g of $M_5$ seeds. 12 seeds were pooled and pulverized using a steel ball and a TissueLyzer bead beater (Qiagen).

For the methanolic extraction, ~15 mg of seed flour were extracted in 1 ml of extractant (60% methanol, 0.06% formic acid, and 15 ppm caffeine in water) for 3 hours at 1200 rpm. The extracts were cleared by centrifugation and diluted 15x with ultrapure water. After filtration through a 0.22-µm filter, the extracts were analyzed by LC-MS as described above under *Analytical methods*. Sparteine was quantified using an external standard curve, with caffeine used as internal standard.

For the acid-base extraction, 250 mg of seed flour were extracted in 2 ml of 1 M $HCl_{(aq)}$ by shaking at 400 rpm for 1 hour at room temperature. Debris was removed by centrifugation and the cleared acidic extract was defatted with n-hexane (5x 1-ml portions). The defatted extract was basified with 1.5 ml of 2 M $NaOH_{(aq)}$ and extracted with dichloromethane (5x 1-ml portions). The organic extract was dried over anhydrous $Na_2SO_4$ and evaporated under reduced pressure. The residual oil was redissolved in 500 µl of ethyl acetate. For determination of the enantiomeric excess, the extract was diluted 5x in ethyl acetate before chiral GC-MS analysis as described under *Analytical methods*.

For the determination of the purity of (–)-sparteine upon acid-base extraction, 15 µl of the ethyl acetate extract were evaporated by gentle heating and the residue was redissolved in 50 µl of 60% aqueous methanol containing 0.06% formic acid. The sample was then diluted 15x with ultrapure water, filtered through a 0.22-µm filter, and analyzed by LC-MS as described above under *Analytical methods*. The purity of the extracted sparteine was estimated using the %area normalization method. All chromatographic peaks visible in the total ion chromatogram of the extract but absent in that of a blank extraction sample were included in the calculation of the total peak area.

*Isolation and chiral analysis of (–)-sparteine from $M_6$ $CYP71D189^{KO}$ seeds*

300 g of $M_5$ seeds derived from forty-four $M_4$ $CYP71D189^{KO}$ homozygous plants were grown in the field in Christchurch, New Zealand. The seeds were machine-sown in a single plot in November 2022. 3 kg of $M_6$ seeds were harvested in April 2023. The seeds were pooled, and a 100 g portion was pulverized using a steel ball and a TissueLyzer bead beater (Qiagen).

10 g of seed flour were extracted in 200 ml of 0.5 M $H_2SO_{4(aq)}$ by stirring at 600 rpm for 18 hours at room temperature. Large debris was removed by centrifugation. The yellow oil layer that formed above the extract was discarded by pipetting. The extract was basified with 35 ml of 10 M $NaOH_{(aq)}$ and centrifuged again to remove the precipitate that formed upon addition of the base. The basified extract was extracted with diethyl ether (5x 50-ml portions). The organic extract was dried over anhydrous $Na_2SO_4$ and evaporated under reduced pressure. The residual oil was redissolved in 2 ml of isopropanol. 2 ml of 0.5 M $H_2SO_4$ in isopropanol were then added to precipitate (–)-sparteine as its bisulfate salt. The mixture was kept for 12 h at 4°C, and the resulting crystals were collected by vacuum filtration, washed with ice-cold isopropanol, and dried over

the filter. (–)-Sparteine bisulfate ($C_{15}H_{26}N_2 \cdot 2H_2SO_4$; 55-71 mg, 0.30-0.39% of seed dry weight) appeared as a white, crystalline powder which decomposed upon heating to 262-264°C (m.p. lit.[8] 247-250°C, decomp.).

To determine the enantiomeric excess and the purity of the isolated (–)-sparteine bisulfate by GC-MS, samples were prepared by dissolving 3 mg of (–)-sparteine bisulfate or commercial (–)-sparteine in 1 ml of 5 mM $H_2SO_{4(aq)}$. 500 µl of the sparteine solutions were basified with 50 µl 10 M $NaOH_{(aq)}$ and extracted with 2 ml ethyl acetate. The organic extract was dried over anhydrous $Na_2SO_4$, diluted 5x in ethyl acetate, and analyzed as described above under *Analytical methods*.
