## Supplementary Table 1 for "Metabolic engineering of narrow-leafed lupin for the production of enantiomerically pure (‒)-sparteine"

**Supplementary Table 1 | Candidate genes for the oxidation of sparteine to lupanine in NLL.** Organ-specific expression profile of the selected oxidases in a bitter NLL cultivar (Oskar) and a sweet cultivar (Tanjil).

| Gene ID from the NLL reference genome (NCBI) | Name abbreviation | Expression level in Tanjil (TPM) | | | | | Expression level in Oskar (TPM) | | | | | | | | Pearson's correlation coefficient (compared to LDC) |
|---|---|---|---|---|---|---|---|---|---|---|---|---|---|---|---|
| | | Root | Stem | Leaf | Flower | Seed | Root | Stem | Leaf | Pedicel | Flower | Mature pericarp | Seed | Young pod | |
| LOC109327937 | LDC | 0.1 | 4.6 | 0.0 | 0.0 | 0.0 | 17.9 | 233.6 | 319.0 | 202.6 | 0.0 | 32.3 | 0.0 | 23.8 | 1 |
| LOC109338642 | CYP76E36 | 15.3 | 6.7 | 0.0 | 4.3 | 0.5 | 9.2 | 38.2 | 50.5 | 27.6 | 0.0 | 13.8 | 3.3 | 8.1 | 0.96 |
| LOC109360201 | CYP71D186 | 1.8 | 1.6 | 0.4 | 2.7 | 0.2 | 13.7 | 45.0 | 63.1 | 52.4 | 0.0 | 27.8 | 0.0 | 13.2 | 0.95 |
| LOC109357725 | CYP71A168 | 0.0 | 11.5 | 1.6 | 1.1 | 0.1 | 4.5 | 96.4 | 167.8 | 34.7 | 0.0 | 44.3 | 0.2 | 21.2 | 0.92 |
| LOC109337773 | SDR1 | 26.7 | 5.0 | 0.0 | 7.6 | 17.7 | 6.8 | 74.1 | 89.3 | 150.7 | 23.4 | 87.1 | 7.4 | 97.3 | 0.68 |
