## Supplementary Table 2 for "Metabolic engineering of narrow-leafed lupin for the production of enantiomerically pure (‒)-sparteine"

**Supplementary Table 2 | List of DNA oligos used in this study.**

| Name | Sequence | Target | Purpose |
|---|---|---|---|
| mGFP5_pEAQ_FW | GGCTTAAUATGAGTAAAGGAGAAGAACTTTTC | *mGFP5* from the plasmid pCAMBIA1302 | USER cloning into pEAQ-USER |
| mGFP5_pEAQ_RV | GGTTTAAUTTATTTGTATAGTTCATCCATGCC | *mGFP5* from the plasmid pCAMBIA1302 | USER cloning into pEAQ-USER |
| CYP71D189_pEAQ_FW | GGCTTAAUATGGAGCTTCAAAACCCTTTC | *CYP71D189* from NLL *cultivar* Oskar | USER cloning into pEAQ-USER and genotyping of $CYP71D189^{KO}$ NLL plants |
| CYP71D189_pEAQ_RV | GGTTTAAUTTAAGGCATACGAGTAACAATTGGA | *CYP71D189* from NLL *cultivar* Oskar | USER cloning into pEAQ-USER and genotyping of $CYP71D189^{KO}$ NLL plants |
| SDR1_pEAQ_U_FW | GGCTTAAUATGGTAGAAACTACTTCCAACAACAG | *SDR1* from NLL *cultivar* Oskar | USER cloning into pEAQ-USER |
| SDR1_pEAQ_U_RV | GGTTTAAUTCATGGAGCAATATAGGAACCATCC | *SDR1* from NLL *cultivar* Oskar | USER cloning into pEAQ-USER |
| CYP76E36_pEAQ_FW | GGCTTAAUATGGATTATCTAACACTTTTTCTACTCA | *CYP76E36* from NLL *cultivar* Oskar | USER cloning into pEAQ-USER |
| CYP76E36_pEAQ_RV | GGTTTAAUTTATGCTTTGATAGGAATAACTAGGAGA | *CYP76E36* from NLL *cultivar* Oskar | USER cloning into pEAQ-USER |
| CYP71A168_pEAQ_FW | GGCTTAAUATGCCTCTTCTATCTATTCAAGAG | *CYP71A168* from NLL *cultivar* Oskar | USER cloning into pEAQ-USER |
| CYP71A168_pEAQ_RV | GGTTTAAUTCAAGGCTGAGATCCACTTC | *CYP71A168* from NLL *cultivar* Oskar | USER cloning into pEAQ-USER |
| CYP71D189_TaqMan_FW | ATATCAGCAATTGAGGAAGG | *CYP71D189* from NLL *cultivar* Oskar | Primer for TaqMan assay for genotyping the mutant NLL DNA library |
| CYP71D189_TaqMan_RV | TCAAGTTTAGCCTTTGTCTT | *CYP71D189* from NLL *cultivar* Oskar | Primer for TaqMan assay for genotyping the mutant NLL DNA library |
| CYP71D189WT_TaqMan_HEX | CAGGAGAACTATGGGTTAGT | $\mathit{CYP71D189}^{WT}$ allele from NLL *cultivar* Oskar | $\mathit{CYP71D189}^{WT}$ allele-specific probe for TaqMan assay containing a HEX fluorophore and a BHQ1 quencer |
| CYP71D189KO_TaqMan_FAM | AGGAGAACTATGAGTTAGTGA | $\mathit{CYP71D189}^{KO}$ allele from NLL *cultivar* Oskar | $\mathit{CYP71D189}^{KO}$ allele-specific probe for TaqMan assay containing a FAM fluorophore and a BHQ1 quencer |
| pEAQ_Seq_FW | GCTTCTGTATATTCTGCCCAAATTCG | pEAQ-USER | Sanger sequencing of pEAQ-USER constructs and culture PCR |
| pEAQ_Seq_RV | CCGCTCACCAAACATAGAAATGC | pEAQ-USER | Sanger sequencing of pEAQ-USER constructs and culture PCR |
